# Associations between multivariate profiles, lifestyle behaviors and reaction time performance in active young adults

**DOI:** 10.64898/2026.10.02.753577

**Authors:** Gloria Valdivia-Tapia, Patricia Huerta, Pablo Vergara-Barra, Jorge Fuentealba, Patricio A. Castro

**Author notes:** Correspondence: Patricio A. Castro.

## Abstract

Lifestyle behaviors influence cardiometabolic and cognitive health through multiple interconnected processes, involving metabolic, cardiovascular, behavioral and cognitive factors. This study aimed to assess associations between multivariate plasma and physiological profiles, and behavioral parameters, and their potential association with reaction time (RT) in active young adults. Data on physical activity levels were collected from wearable devices and GPAQ questionnaire, along with anthropometric and clinical assessments, cognitive tests, and blood analyses. Principal component analysis was performed separately for physiological and plasma parameters, retaining the first two principal components (PCs) from each analysis. Multiple linear regression models were used to evaluate the associations between plasma, sex, physical activity levels (METs), daily step count, and alcohol/marijuana use on physiological PC1/PC2 scores, as well as RT. Continuous independent variables were previously categorized, and combined variables were created to represent specific combinations between plasma/physiological PC1 categories and sex. Physiological PC1 (blood pressure) was associated with combined plasma PC1 (insulin/HOMA-IR)-sex variable, total METs and alcohol consumption (adjusted R^2^ = 0.30). Physiological PC2 (handgrip strength) was mainly explained by combined plasma PC1-sex variable (adjusted R^2^ = 0.54), with minimal contribution from behavioral factors. RT was associated with physiological PC1 variable and sex (adjusted R^2^ = 0.35), with slower responses observed among women with lower physiological PC1 scores or hypotension; these associations were supported by RT-physiological PC1 and RT-blood pressure correlations. These exploratory findings suggest sex-dependent associations between physiological, metabolic and lifestyle-related factors, supporting a holistic approach in future research, particularly among young adults.

## 1. INTRODUCTION

Lifestyle behaviors, including sedentary behavior and substances use, are important modifiable factors influencing health. Globally, about 1.8 billion adults did not meet recommended levels of physical activity (PA) in 2022, while in Chile, approximately 31.5% of the adult population was classified as physically inactive according to WHO recommendations (Fernández-Verdejo and Suárez-Reyes, 2021; Strain et al., 2024). In addition, insufficient PA and sedentary behavior may coexist with other health- related behaviors, including substance use, particularly among adolescents and young adults (West et al., 2020; Santos et al., 2026). Together, they represent modifiable aspects of lifestyle that may be associated with multiple physiological systems.

Healthy lifestyle behaviors, including regular PA, are related with favorable metabolic and cardiovascular (CV) profiles, and may contribute to prevention of cardiometabolic diseases, including type 2 diabetes mellitus, myocardial infarction and stroke (Eroglu et al., 2024). These associations may be reflected in modifications on lipid profiles – as increased HDL cholesterol (HDL-c), and decreased LDL cholesterol (LDL-c) and triglycerides (TG)–, as well as improved insulin sensitivity, commonly estimated using HOMA-IR (Franczyk et al., 2023). Multifactorial lifestyle interventions are an effective strategy for reducing CV risk factors, including blood pressure (BP), lipids, body mass index (BMI) and waist circumference (Sisti et al., 2018). In contrast, alcohol use has been associated with alterations in CV parameters, as BP and heart rate (HR), while marijuana use has also been related to acute alterations in CV function (Pacher et al., 2018; Arora et al., 2022). These factors can also influence the function and regulation of the autonomic nervous system, resulting in changes in resting HR, through the modulation of sympathetic and parasympathetic activity (Fu and Levine, 2013; Tiwari et al., 2021). Combined, these relationships reflect the interconnected nature of metabolic, CV and autonomic systems, and their response to external and modifiable lifestyle factors.

This associations extends to the central nervous system, where CV, metabolic, autonomic, and neural interactions contribute to brain functions. Related to substance consumption, frequent marijuana uses and higher levels of alcohol use have been associated with impairment in cognitive performance, although these effects may vary according to dose and exposure patterns (Lees et al., 2021; Weinstock et al., 2021). In contrast, regular PA and exercise have been related with neuroplastic processes and structural changes in areas as hippocampus and cortex, which are involved in cognitive processes, such as memory, learning, and attention (Erickson et al., 2011; Kandola et al., 2016; Esteban-Cornejo et al., 2021). Actually, several studies support associations between PA, exercise, and reaction time (RT), which involves attention, as well as premotor, motor and processing functions(Cech and Martin, 2012; Reigal et al., 2019). This evidence suggest that physical, mental and cognitive health are interconnected and contribute to overall well-being. Grip strength, for example, is an objective measure of muscular fitness, that has been evaluated as a reliable predictor of cognitive health (Firth et al., 2018).

Consequently, considering the interplay among these systems may provide a more comprehensive understanding of factors associated with physiological profiles and cognitive performance. This preliminary study aimed to integrating and evaluated associations of plasma and behavior variables – PA levels, alcohol/marijuana use– and sex, as potential parameters that contribute to these physiological profiles and reaction time performance in active young adults, using principal component analysis and multiple linear regression models.

## 2. MATERIALS AND METHODS

### 2.1. EXPERIMENTAL DESIGN & PARTICIPANTS

This observational and exploratory study collected data on lifestyle behaviors among young adults, between September 2022 and April 2023. Target population consisted of undergraduate students aged 19 to 27 years, who were invited to participate in the study. A simple random sampling was performed among those subjects who responded to the call and the final sample consisted only of those who completed all assessments and provided all the required information. Participants with missing data were excluded. In a single online session, all participants were provided with an informed consent approved by the Ethics Committee of the pharmacy school of the university, and a sociodemographic and the Global Physical Activity Questionnaire (GPAQ) to complete (World Health Organization, 2012).

### 2.2. BEHAVIOURAL, PHYSIOLOGICAL & COGNITIVE PARAMETERS

Sociodemographic questionnaire collected information on age, gender and consumption habits (alcohol, tobacco and marijuana), using 5 points Likert scales –based on AUDIT questionnaire (Babor et al., 2001). PA levels were expressed in MET, where 1 MET is equivalent to basal energy expenditure at rest (1 kcal kg^-1^ hour^-1^). Total METs were calculated from GPAQ results, as the sum of METs from vigorous PA (vigorous METs) and moderate PA (moderate METs) (World Health Organization, 2012). PA levels were categorized according to total METs volume in MET-hours per week (MET-h/wk): inactive (<3.5), low (3.75-7.49), moderate (7.5-16.49), high (16.50-25.49) and very high (≥25.50) (Chang et al., 2020). Each participant received a wearable device (Fitbit smartband) to record their daily step count and resting HR (bpm) as indicators of regular PA (Castro et al., 2022). After 3 weeks of free use, each participant entered the data recorded into an online form, hosted on the institutional web platform. Records from seven different days were requested.

Then, participants were scheduled to attend sessions for physiological measurements and cognitive tests. Systolic and diastolic BP (mmHg) were measured twice using a wrist monitor GM-500 W (Generation Guard). BMI (kg/m^2^) was calculated from body weight, measured using ESF14 Smart Fitness Scale (Etekcity), and height, measured using a tape measure. Handgrip strength (kg) was determined twice on each hand, using an analog hand dynamometer (Jamar), according to procedures described by Firth et al. (Firth et al., 2018).

Cognitive performance was assessed using RT and visuospatial memory tests from the Human Benchmark web-based platform (Cuthbertson et al., 2015). All tests were performed under standardized conditions using the same equipment. The RT test recorded the response time (milliseconds, ms) to the color change from red to green on the screen and the score for each participant was defined as the mean response time across five trials. In the visuospatial memory test, participants had to recall increasingly complex patterns of squares on a grid and select them after they disappeared; the highest level achieved by each participant was used as the score.

### 2.3. PLASMA PARAMETERS

For the analysis of plasma parameters, blood samples were collected from volunteers after an overnight fast (∼10-12 hours) at a certified laboratory (Institute of Public Health [ISP], Chile). Lipid profiles (total cholesterol [TC], high-density lipoprotein cholesterol [HDL-C], triglycerides [TG]; mg/dL), liver enzymes [aspartate aminotransferase [AST] and alanine aminotransferase [ALT], U/L), and blood glucose (mg/dL) levels were determined using colorimetric methods. Insulin (uIU/L) and high-sensitivity C reactive protein (CRP, mg/L) were measured by ELISA. Low-density lipoprotein cholesterol (LDL-C, mg/dL) was calculated using the Friedewald equation (Friedewald et al., 1972), and insulin resistance was estimated by the homeostatic model assessment (HOMA-IR) according to the method of Matthews et al. (Matthews et al., 1985).

### 2.4. STATISTICAL ANALYSIS

Continuous variables were expressed as mean ± standard error of the mean (SEM). Data distribution was assessed using the Shapiro-Wilk test, and comparisons between sexes were performed using the Student’s t-test or the Mann-Whitney U test, as appropriate. Statistical significance was defined by p-values < 0.05, with 95% confidence interval, in all analyses.

Correlation analyses between plasma and physiological variables was performed using Spearman’s coefficient, to observe associations of medium to high strength (R > 0.5) and significance <0.05. To reduce dimensionality of the original variables, principal component analyses (PCA) were performed separately on ten plasma parameters –previously described– and five physiological variables: resting HR, systolic and diastolic BP, handgrip strength, and BMI. PCA specifications included a minimum of 4 observations per variable, no missing values, the assumption of linearity, correlation among variables –previously checked with Spearman’s R–, and identification and exclusion of outliers. PCs selection was based on the cumulatively explained variance (eigenvalues and elbow of the scree plot), and were used in subsequent analyses. In both cases, the first two principal components (PCs) were retained and designated as “plasma PC1/PC2” and “physio PC1/PC2”.

Then, multiple linear regression models were developed with physio PC1 and PC2, and RT as continuous dependent variables. As independent variables, plasma and physio PC1/PC2 scores, sex and all behavioral variables –total METs, vigorous METs, daily steps count, alcohol and marijuana use– were categorized. Plasma and physiological PC1/PC2 scores were categorized into deciles and grouped into low (1-3), medium (4-6) and high (7-10) levels. Total and vigorous METs were similarly classified into deciles and grouped into low (1-3) and high (4-10) levels, while daily step count was categorized as low (<10000 steps/day) or high (≥10000 steps/day). Levels of substances used were defined according to frequency of use as low (0-2) and high (3-4). Initially, the association of each independent variable with the dependent variables was assessed, followed by the progressive incorporation of additional variables into the models, in order to evaluate their individual effects and potential interactions, and evaluate their potential confounding role –assessed as percentage of change in coefficients. Interactions between PCs categories and sex were recoded into a four-level combined variable: low PC-women, high PC (medium and high)-women, low PC (low and medium)-men and high PC-men. Models were compared based on the coefficient of determination (R²) and adjusted R². Comparisons between models were performed using the Akaike Information Criterion (AIC) and F Statistic, residual analyses were subsequently performed to assess models and to verify its assumptions.

## 3. RESULTS

### 3.1. Characterization of the study population

This study included a total of 30 participants: 20 men and 10 women. The variables assessed, as described in **Table 1**, were classified into four categories: behavioral, physiological, cognitive, and plasma parameters. Concerning behavioral parameters, very high levels of PA (≥ 25.50 MET-h/week) were mainly reported and no values associated to low or inactive levels (≤ 7.49 MET-h/week) were recorded among the students (Chang et al., 2020). Alcohol and marijuana use were more widespread than tobacco use. Physiological differences were observed between men and women in resting HR (p= 0.0343) and handgrip strength (p < 0.0001). The mean BMI was in “normal weight” range for women (24.6 kg/m^2^), and within the “overweight” category for men (25.1 kg/m^2^), according to WHO BMI categories (World Health Organization, 2000). Moreover, men had faster RT than women (p = 0.0193), while no statistically significant differences by sex were observed in the visuospatial memory test. Related to plasma parameters, there were differences by gender in glucose (p = 0.0477), insulin (p = 0.0079), HOMA-IR (p = 0.0189), AST (p = 0.0484) and CRP (p = 0.0412, n = 28).

**Table 1.** Characteristics of Study Subjects.

|  | OVERALL | MEN | WOMEN |
| --- | --- | --- | --- |
| n | 30 | 20 | 10 |
| Age (years) | 22 (0.38) | 22 (0.46) | 21 (0.63) |
| **BEHAVIORAL PARAMETERS** |  |  |  |
| Total METs (-h/week) | 64 (8.1) | 58 (6.4) | 78 (21) |
| Vigorous METs (-h/week) | 31 (6.9) | 30 (6.2) | 35 (17) |
| Daily Steps (No.) | 10639 (709) | 11115 (747) | 9687 (1532) |
| Substances Consumption (%) <sup>a</sup> |  |  |  |
| Alcohol | 93.3 | 90.0 | 100.0 |
| Marijuana | 36.7 | 45.0 | 20.0 |
| Tobacco | 6.7 | 10.0 | 0.0 |
| **PHYSIOLOGICAL PARAMETERS** |  |  |  |
| Resting Heart Rate (bpm) <sup>b</sup> | 64 (1.5) | 62 (1.7) | 69 (2.5) |
| Systolic Blood Pressure (mmHg) | 119 (3.1) | 121 (3.9) | 114 (4.7) |
| Diastolic Blood Pressure (mmHg) | 74 (1.8) | 75 (2.0) | 71 (3.3) |
| Handgrip (kg) <sup>b</sup> | 34 (2.1) | 40 (2.1) | 23 (1.4) |
| Body Mass Index (kg/m<sup>2</sup>) | 24 (0.59) | 24 (0.67) | 25 (1.2) |
| **COGNITIVE PARAMETERS** |  |  |  |
| Reaction Time (ms) <sup>b</sup> | 272 (6.3) | 261 (4.9) | 294 (14) |
| Visuospatial Memory (level) | 10 (0.36) | 10 (0.44) | 9.9 (0.62) |
| **PLASMA PARAMETERS** |  |  |  |
| Total Cholesterol (mg/dL) | 153 (5.5) | 153 (6.8) | 153 (9.8) |
| HDL-C (mg/dL) | 57 (2.4) | 55 (2.7) | 62 (4.8) |
| LDL-C (mg/dL) | 92 (8.1) | 91 (9.8) | 95 (15) |
| Triglycerides (mg/dL) | 74 (4.5) | 77 (5.6) | 68 (7.7) |
| Glucose (mg/dL) <sup>c</sup> | 93 (0.95) | 95 (1.2) | 91 (1.4) |
| Insulin (μUI/mL) <sup>b</sup> | 10 (0.83) | 8.6 (0.58) | 13 (1.9) |
| HOMA-IR <sup>b</sup> | 2.3 (0.20) | 2 (0.15) | 3 (0.47) |
| GOT (U/L) <sup>b</sup> | 19 (1.1) | 21 (1.4) | 16 (1.2) |
| GPT (U/L) | 18 (1.1) | 19 (1.4) | 15 (1.4) |
| C Reactive Protein (mg/L) <sup>c, d</sup> | 4.7 (3.3) | 6.5 (5.0) | 1.2 (0.29) |

Values are expressed as Mean (SEM). <sup>a</sup> Percentages (%) associated to frequency: recurrent (≥ 4 times/week), regular (2-3 times/week), occasional (2-4 times a month) and rarely (≤1 time/month). Difference of average by gender (p value <0.05): <sup>b</sup> t-test (Welch's correction) or <sup>c</sup> Mann-Whitney test. <sup>d</sup> Participants with values ≥ 10 mg/L were excluded for descriptive analysis (men, n=18).

### 3.2. Principal component analyses

PCA of plasma variables (**Figure 1A**) showed that plasma PC1 and PC2 accounted for 45.0% of the total variance –25.3% and 19.7%, respectively. The main contributors to plasma PC1 were insulin and HOMA-IR (31.5% and 30.2%, respectively), indicating that this component was primarily characterized by a metabolic profile related to insulin resistance. Furthermore, TC and LDL-C were the main contributors to plasma PC2 (29.0% and 22.8%) followed by AST (18.5%), suggesting a lipid-related metabolism axis with a hepatic component (**Figure 1B**). PCA of physiological variables, showed that PC1 and PC2 accounted for 73.1% of the total variance –45.2% and 27.9%, respectively (**Figure 1A**). Physio PC1 was primarily characterized by systolic and diastolic blood pressure, which contributed 39.3% and 38.5%, respectively, indicating a cardiovascular/hemodynamic profile. In contrast, the main contributors of PC2 handgrip strength (54.0%), followed by resting HR (44.6%) (**Figure 1C**), suggesting a physical fitness-related profile.

**Figure 1.**
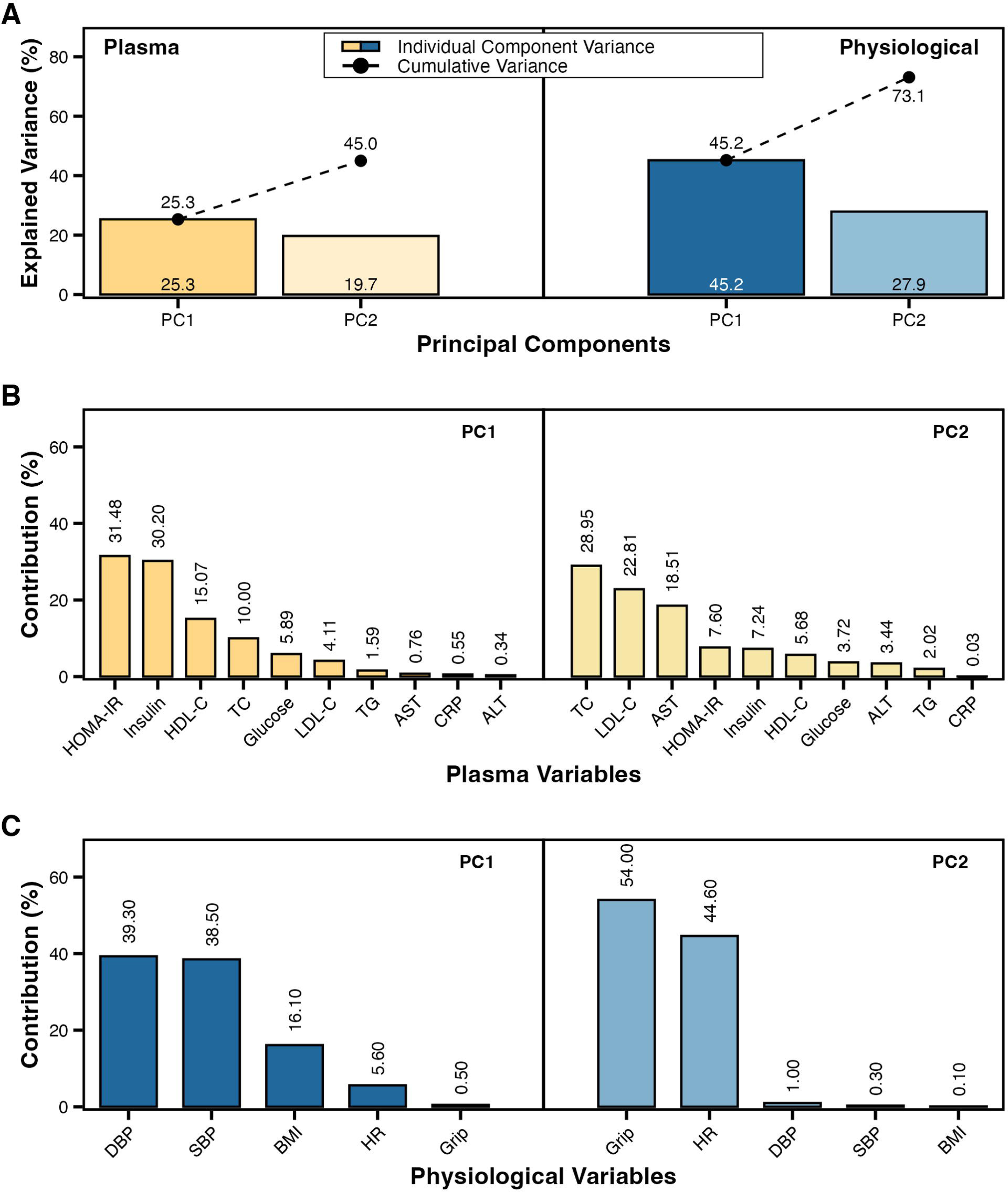
Principal Component Analysis, PCA. (**A**) Variance explained (%) by each principal component (PC; bars) and cumulative variance (connected dots) for physiological and plasma analyses (panels). (**B-C**) Relative contribution (%) of each variable to PC1 and PC2, shown separately for (**B**) physiological and (**C**) plasma analyses. *TC*: Total Cholesterol, *TG*: Triglycerides. *DBP*: Diastolic Blood Pressure. *SBP*: Systolic Blood Pressure. *HR*: Resting Heart Rate.

### 3.3. Multiple linear regression models: physio PCs

The first model, with physio PC1 as the dependent variable, showed the best initial fit with the combined plasma PC1-sex variable (baseline model), with low plasma PC1-women as the reference category (**Supplementary Table**). After incorporating total METs, changes were observed in the coefficients across all plasma PC1-sex categories, from 7.9% to 43.8% relative to the baseline model. R^2^ and adjusted R^2^ increased from 0.26 to 0.31 and from 0.18 to 0.20, respectively. Then, the inclusion of alcohol consumption also resulted in changes in the plasma PC1-sex coefficients (up to 13%) and in total METs coefficients (35%). In this final model, the coefficients for women and men with high plasma PC1 were significant (p < 0.05 and p < 0.01, respectively). Also, it showed a lower AIC than the previous model (107.32 vs. 110.40), with an R^2^ of 0.42 and adjusted R^2^ of 0.30. These results suggest that physio PC1 –BP– was associated with the combined plasma PC1-sex variable. In addition, it was positively associated with PA and inversely associated with alcohol use (**Figure 2A**).

**Figure 2:**
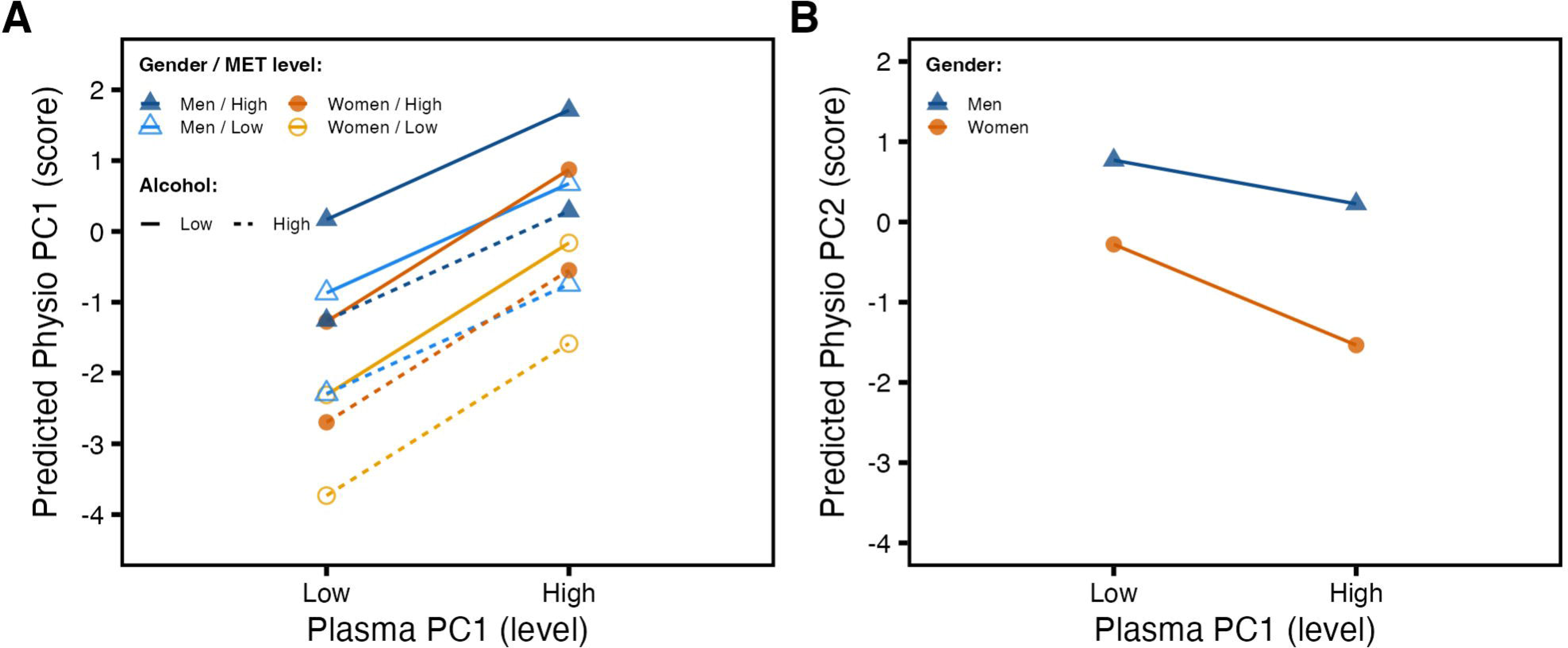
Multiple Linear Regression Model of Physiological PCs. Predicted values from fitted models; both are based on the combined variable plasma PC1-sex, defined by specific combinations of plasma PC1 levels and sex. (**A**) Model for physio PC1 adjusted for total METs and alcohol consumption level. (**B**) Model for physio PC2.

The second model, with physio PC2 as the dependent variable, also showed the best initial fit with the combined plasma PC1-sex variable (**Supplementary Table**). This model showed an R^2^ of 0.59 and an adjusted R^2^ of 0.54. Unlike the physio PC1 model, inclusion of any additional behavioral variables did not improve the model fit. Although addition of alcohol consumption slightly increased R^2^ to 0.61, neither adjusted R^2^ nor AIC indicated an improvement in fit. Therefore, the initial model was retained for physio PC2, which was primarily characterized by handgrip strength (**Figure 2B**).

### 3.4. Reaction time

Based on RT, potential associations with plasma, physiological and behavioral parameters were assessed. Among the analyses performed, the multiple linear regression model including physio PC1 and sex as independent variables showed the best fit for RT (R^2^ = 0.42 and adjusted R^2^ = 0.35, **Figure 3A**) than models that included only physio PC1 (R^2^ = 0.32, adjusted R^2^ = 0.27) or only sex (R^2^ = 0.21, adjusted R^2^ = 0.18). Women and low physio PC1 were used as reference categories for sex and each categorical independent variable, respectively. Inclusion of physio PC2, plasma PC1 or PC2, or behavioral factors did not improve model fit, while a model with a combined variable failed the homoscedasticity assumption (**Supplementary Table**). In contrast, models for visuospatial memory did not provide optimal fits.

**Figure 3:**
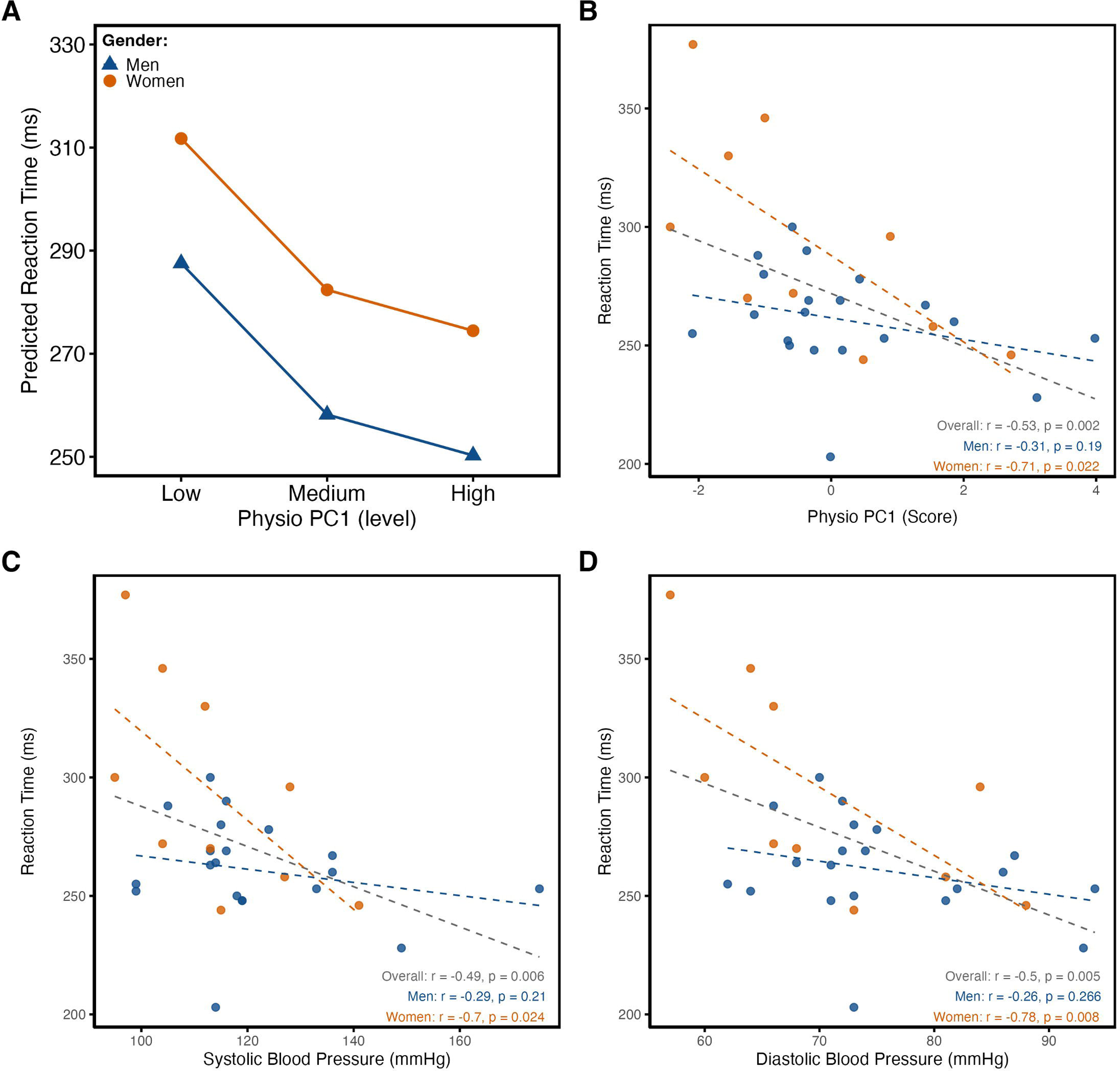
Reaction time. (**A**) Predicted reaction time from the multiple linear regression model including the physio PC1 and sex variables. (**B-D**) Correlation plots of reaction time with (**B**) physio PC1 scores, (**C**) systolic BP and (**D**) diastolic BP. Spearman correlation coefficients (R) and p-values are shown for men (blue), women (orange), and the overall sample.

Then, analysis of predicted mean RT revealed lower RT among men –in any physio PC1 category–and women with high PC1 level, compared to reference category –women with low physio PC1– (**Figure 3A**). Furthermore, the relationship between observed RT and physio PC1 scores showed a different distribution profile among women, with a statistically significant inverse correlation (R = −0.71, p < 0.05; **Figure 3B**). In contrast, the distribution pattern observed in men was more homogeneous, with no significant correlation between RT and physio PC1 scores (R = −0.31, p>0.05). A similar inverse association pattern was observed between RT and BP, with significant correlations for both systolic (R = −0.70, p<0.05; **Figure 3C**) and diastolic BP (R = −0.78, p<0.05; **Figure 3D**). Overall, these findings indicate that lower physio PC1 levels, mainly characterized by lower BP, were related with longer RT, with a strong association between physio PC1 and RT particularly evident among women.

## 4. DISCUSSION

The study population was mainly characterized by being active, with high or very high levels of PA, according to estimated total METs. These findings may be related to the high daily step count (mean: >10000 steps/day), promoted by the large university campus, which supports active walking among students. Evidence suggests that, rather than a specific amount, any type of PA provides health benefits, especially in the context of actual lifestyles, which encourage inactivity and sedentary behaviors (Piercy et al., 2018). Concerning unhealthy habits, alcohol consumption exceeded marijuana and tobacco, independent of frequency (93.3%, 36.7% and 6.7%, respectively). Previous report has identified alcohol and marijuana as commonly used substances among Chilean undergraduate students; although their use was declined between 2021-2025, its frequency or intensity remains a cause for concern (Servicio Nacional para la Prevención y Rehabilitación del Consumo de Drogas y Alcohol, 2025). Despite the coexistence of both –healthy and unhealthy habits–, this group of young adults reported no history of CV diseases or impaired biochemical parameters, except for CRP in men (outsiders, excluded only from descriptive analyses) and HOMA-IR in women. Even so, if abnormalities were present, they might be subtle and not necessarily apparent when analyzing parameters individually.

PCA allowed us to identify multivariate plasma and physiological profiles by integrating information from several correlated variables into common dimensions. Previous studies have applied PCA to cardiometabolic risk factors in a single analysis, incorporating metabolic, CV, anthropometric and behavioral variables (Peterson et al., 2012). In contrast, given the objective of this study, plasma and physiological variables were analyzed independently, allowing to identify specific multivariate dimensions for each domain. Plasma PCA showed two different metabolic profiles within the study population (**Figure 1B**). PC1 was predominantly characterized by insulin and HOMA-IR, suggesting an insulin resistance related dimension, while PC2 was characterized mainly by total cholesterol and LDL-C, with a lower contribution from AST, indicating a lipid-related dimension. In contrast, physiological PCA identified a CV/hemodynamic physio PC1, mainly characterized by systolic and diastolic BP, and a physio PC2 reflecting a muscular fitness axis, characterized by handgrip strength (**Figure 1C**). These PCA-derived dimensions may reveal less favorable profiles, despite individual values remained within clinically accepted reference ranges (Burrows et al., 2015; Jones et al., 2025), and were incorporated into regression models to assess associations between plasma and physiological profiles, as well as the influence of behavioral factors.

For both physiological components, the final models retained the combined plasma PC1-sex variable, but not plasma PC2, suggesting that the metabolic profile, predominantly characterized by insulin and HOMA-IR, was more closely associated with the physiological profiles assessed in this study. This finding is consistent with the reported association between insulin resistance, assessed by HOMA-IR, and blood pressure (Quesada et al., 2021). Also, the association between plasma PC1 and the physiological components was modified by sex, as reflected by the combined plasma PC1-sex variable in final models, although the definition of this variable limits the direct comparison of its categories between men and women, as the corresponding categories represented different ranges of the PC1 distribution. Regarding the physio PC1 model, representing the CV dimension characterized mainly by BP, total METs and alcohol consumption were retained in the model (**Figure 2A**), which was statistically significant (**Supplementary Table**). Inclusion of total METs is consistent with the established relationship to physical activity, insulin sensitivity and cardiometabolic health (U.S. Department of Health and Human Services, 2018). Other behavioral parameters, such as vigorous METs, daily step count and marijuana use, did not improve the model fit. Therefore, these preliminary models suggest that behavioral, metabolic and CV factors may no act independently, but rather converge within interconnected physiological profiles.

To assess if these physiological profiles identified by PCA were also reflected in cognitive performance, we evaluated reaction time and visuospatial memory. The final model for RT (F = 6.16, p = 0.0026) explained 35.0% of the variance (adjusted R^2^), with physio PC1 and sex as the only parameters retained (**Figure 3A**); no significant regression model was identified for visuospatial memory. In contrast to previous models of physio PCs, in which specific categorization by sex limited direct comparisons, analysis of RTs predicted by this model showed differences by sex. Although RT is considered a relatively simple measure, defined as the interval between stimulus presentation and motor response, central processing accounts for approximately 80% of total RT (Cech and Martin, 2012). RT also involves processes such as attention, concentration, cortical arousal and processing speed (Cech and Martin, 2012; Prabu Kumar et al., 2020). Faster visual RT have been associated with better cognitive function and academic performance, with Prabu Kumar et al. (2020) reporting faster RTs among predictors of better academic performance and that female students exhibited faster RTs than male. Therefore, in addition to sex, this model suggests that the physiological dimension previously identified may also be associated with cognitive-motor performance.

Supporting the regression results, correlation analyses suggest that the association between the physiological profiles (physio PC1) and RT was closely related to its blood pressure components. This relationship was more evident among women than men, with relatively strong inverse correlations observed, despite the smaller subgroup of women (n=10 vs. n=20). Women with lower systolic BP (<100 mmHg) and lower diastolic BP, as well as lower physio PC1 values, showed longer RTs, contributing to the overall inverse association observed in this group (**Figure 3B-3D**). This pattern was less pronounced among men, with very weak inverse correlations. Although research on CV risk and cognitive performance, including RT and memory, has mainly focused on elevated BP and hypertension, lower BP has also been associated with poorer cognitive performance (Duschek et al., 2003). Duschek et al. (2003) reported reduced attentional performance and prolonged RTs in individuals with moderate hypotension, suggesting that lower BP may be associated with impaired cognitive performance, potentially related to altered cerebral perfusion and reduced metabolic rate, altered baroreceptor activity, and reduced well-being. Also, these authors hypothesized an inverted U-shaped relationship between BP and cognitive function, with a presumably optimal performance at normotensive levels. The inverse associations observed in this study between BP and RT, particularly among women, may be consistent with the lower-BP section of a non-linear relationship. However, the data do not allow to established this association, mainly because we do not observed evidence of higher BP levels.

Finally, this preliminary study suggests that metabolic, physiological and behavioral parameters may be interconnected with cognitive performance, represented by RT, instead of acting as separate domains. This integrated perspective is consistent with the American Heart Association’s “Life’s Essential 8”, which incorporates behavioral and health factors as key components of CV health (Lloyd-Jones et al., 2022). Notably, the relationships identified in the present study differed according to sex, particularly in RT with a stronger association observed among women, despite their smaller representation in the study population. In future studies, these models should be evaluated in larger and more balanced groups, potentially revealing associations that were not detected in the current analysis. For example, daily step count did not improve any of the models related to physiology and RT, although higher daily step count has previously been associated with lower resting HR and reduced all-cause mortality (Saint-Maurice et al., 2020; Castro et al., 2022). Furthermore, other lifestyle factors, such as diet and sleep, were not assessed and may represent additional sources of variability in the observed profiles. Similarly, although most participants self-reported as healthy, diagnosed pathologies and pharmacological treatments were not assessed, despite evidence that certain medications, including antidepressants, may affect BP (Calvi et al., 2021). Follow-up of this cohort could further determine whether the multivariate profiles identified are associated with changes in physiological and cognitive performance over time. Given the complex interplay among lifestyle factors, physiological and biochemical characteristics, and cognitive performance, future research should validate these multivariate profiles, particularly in young adults to better understand health trajectories in the context of an increasingly aging population.

## Notes

### Competing Interest Statement

The authors have declared no competing interest.

### Funding

- Ministerio de Educación de Chile: UCO 1895
- Agencia Nacional de Investigación y Desarrollo: FONDECYT 1231038
- Universidad de Concepción: VRIM 2405, VRIM CB2107

