## Supplemental Material for "Associations between multivariate profiles, lifestyle behaviors and reaction time performance in active young adults"

**Supplementary Table 1. Principal Component Analyses (PCA).** Loadings, eigenvalues, percentages of variance and cumulative variance for the first four PCs of (**A**) plasma and (**B**) physiological analyzed variables.

| **A. PLASMA PCA** | **PC 1** | **PC 2** | **PC 3** | **PC 4** |
|---|---|---|---|---|
| TC | -0.503 | 0.756 | 0.327 | -0.006 |
| HDL-C | -0.617 | 0.335 | -0.285 | -0.336 |
| TG | 0.201 | -0.200 | 0.429 | 0.048 |
| LDL-C | -0.322 | 0.671 | 0.478 | 0.228 |
| Glucose | 0.386 | 0.271 | 0.379 | -0.464 |
| Insulin | 0.874 | 0.378 | -0.068 | 0.111 |
| HOMA-IR | 0.892 | 0.387 | -0.035 | 0.050 |
| AST | -0.139 | -0.604 | 0.415 | -0.039 |
| ALT | 0.092 | -0.261 | 0.757 | -0.135 |
| CRP | -0.118 | -0.023 | 0.093 | 0.860 |
| Eigenvalue | 2.527 | 1.973 | 1.504 | 1.157 |
| Variance (%) | 25.3 | 19.7 | 15.0 | 11.6 |
| Cumulative Variance (%) | 25.3 | 45.0 | 60.0 | 71.6 |
| # Variables | | 10 | | |

| **B. PHYSIOLOGICAL PCA** | **PC 1** | **PC 2** | **PC 3** | **PC 4** |
|---|---|---|---|---|
| Resting HR | 0,356 | -0,789 | 0,116 | 0,486 |
| Systolic BP | 0,933 | 0,062 | -0,322 | 0,025 |
| Diastolic BP | 0,943 | 0,117 | -0,236 | -0,144 |
| Handgrip | 0,105 | 0,869 | 0,129 | 0,466 |
| BMI | 0,602 | 0,035 | 0,776 | -0,184 |
| Eigenvalue | 2,261 | 1,396 | 0,791 | 0,508 |
| Variance (%) | 45,2 | 27,9 | 15,8 | 10,2 |
| Cumulative Variance (%) | 45,2 | 73,1 | 89,0 | 99,1 |
| # Variables | | 5 | | |

**Supplementary Table 2.** **Multiple Linear Regression Models Predicting Physio PC1.** Regression coefficients (95% CI) are reported for each category of independent variables, from the base model, with combined variable Plasma PC1-Sex, to the final model. Model fit statistics include $R^2$, adjusted $R^2$, AIC and the F statistic.

| | Models | | |
|---|---|---|---|
| | **Combined (Plasma PC1 - Sex)** | **(Plasma PC1 - Sex) + Total MET** | **(Plasma PC1 - Sex) + Total MET + Alcohol** |
| Intercept | -1.75* (-3.39/-0.1) | -2.51* (-4.54/-0.49) | -2.31* (-4.22/-0.4) |
| High PC1 - Men | 2.76** (0.79/4.72) | 2.98** (1/4.95) | 2.98** (1.13/4.84) |
| Low PC1 - Men | 1.46 (-0.37/3.28) | 1.63 (-0.19/3.46) | 1.44 (-0.28/3.16) |
| High PC1 - Women | 2.03* (0.06/3.99) | 2.47* (0.41/4.53) | 2.15* (0.19/4.11) |
| High METs levels | | 0.77 (-0.44/1.97) | 1.04 (-0.13/2.2) |
| High substance use | | | -1.42* (-2.82/-0.03) |
| $R^2$ / Adjusted $R^2$ | 0.26 / 0.18 | 0.31 / 0.20 | 0.42 / 0.30 |
| AIC | 110.39 | 110.40 | 107.32 |
| F Statistic | 3.12* | 2.83* | 3.46* |

Reference: "low" levels and "women". p-values: (*) <0.05, (**) <0.01, (***) <0.001

**Supplementary Table 3. Multiple Linear Regression Models Predicting Physio PC1 Using Plasma and Sex Variables.** Regression coefficients (95% CI) are reported for each category of independent variables. Model fit statistics include $R^2$, adjusted $R^2$, AIC and the F statistic.

| | **Models** | | | |
|---|---|---|---|---|
| | **Plasma PC1** | **Plasma PC2** | **Sex** | **Plasma PC1 + Sex** |
| Intercept | -1.06* (-1.98/-0.14) | -0.35 (-1.41/0.7) | -0.33 (-1.32/0.67) | -1.49* (-2.65/-0.33) |
| Medium PC1/2 | 0.98 (-0.32/2.29) | 0.88 (-0.61/2.37) | | 0.91 (-0.39/2.21) |
| High PC1/2 | 1.91** (0.69/3.13) | 0.23 (-1.17/1.62) | | 1.97** (0.75/3.18) |
| Men | | | 0.49 (-0.73/1.71) | 0.65 (-0.43/1.73) |
| $R^2$ / Adjusted $R^2$ | 0.28 / 0.22 | 0.06 / -0.01 | 0.02 / -0.01 | 0.32 / 0.24 |
| AIC | 107.82 | 115.87 | 114.89 | 108.1 |
| F Statistic | 5.21* | 0.80 | 0.68 | 4.05* |

Reference: "low" levels and "women". p-values: (*) <0.05, (**) <0.01, (***) <0.001

**Supplementary Table 4. Multiple Linear Regression Models Predicting Physio PC1 Using Behavioral Variables.** Regression coefficients (95% CI) are reported for each category of independent variables. Model fit statistics include $R^2$, adjusted $R^2$, AIC and the F statistic.

| | **Models** | | | | |
|---|---|---|---|---|---|
| | **Daily Steps** | **Total MET** | **Vigorous MET** | **Alcohol** | **Marijuana** |
| Intercept | 0.09 (-0.76/0.94) | -0.21 (-1.27/0.85) | -0.17 (-1.23/0.89) | 0.19 (-0.43/0.80) | 0.05 (-0.57/0.66) |
| > 10000 steps/day | -0.18 (-1.34/0.99) | | | | |
| High METs levels | | 0.30 (-0.96/1.57) | 0.25 (-1.02/1.51) | | |
| High substance use | | | | -1.11 (-2.61/0.39) | -0.45 (-2.38/1.48) |
| $R^2$ / Adjusted $R^2$ | 0.00 / -0.03 | 0.01 / -0.03 | 0.01 / -0.03 | 0.08 / 0.04 | 0.01 / -0.03 |
| AIC | 115.51 | 115.35 | 115.44 | 113.24 | 115.36 |
| F Statistic | 0.10 | 0.24 | 0.16 | 2.30 | 0.23 |

Reference: “<10000 steps/day” and "low" levels. p-values: (*) <0.05, (**) <0.01, (***) <0.001

**Supplementary Table 5. Multiple Linear Regression Models Predicting Physio PC2.** Regression coefficients (95% CI) are reported for each category of the combined plasma PC1-Sex variable. With combined variable Plasma PC1-sex, model fit statistics include $R^2$, adjusted $R^2$, AIC and the F statistic.

| | **Model** |
|---|---|
| | **Combined (Plasma PC1 – Sex)** |
| Intercept | -0.28<br>(-1.24/0.68) |
| High PC1 - Men | 0.50<br>(-0.64/1.65) |
| Low PC1 - Men | 1.05<br>(-0.02/2.12) |
| High PC1 - Women | -1.26*<br>(-2.41/-0.11) |
| $R^2$ / Adjusted $R^2$ | 0.59 / 0.54 |
| AIC | 78.25 |
| F Statistic | 12.58*** |

Reference: "low" levels and "women". p-values: (*) <0.05, (**) <0.01, (***) <0.001

**Supplementary Table 6. Multiple Linear Regression Models Predicting Physio PC2 Using Plasma and Sex Variables.** Regression coefficients (95% CI) are reported for each category of independent variables. Model fit statistics include $R^2$, adjusted $R^2$, AIC and the F statistic.

| | Models | | | |
|---|---|---|---|---|
| | **Plasma PC1** | **Plasma PC2** | **Sex** | **Plasma PC1 + Sex** |
| Intercept | 0.23<br>(-0.53/0.99) | 0.15<br>(-0.69/1.00) | -1.16***<br>(-1.73/-0.59) | -0.84*<br>(-1.54/-0.13) |
| Medium PC1/2 | 0.37<br>(-0.71/1.45) | -0.08<br>(-1.28/1.11) | | 0.19<br>(-0.59/0.98) |
| High PC1/2 | -0.85<br>(-1.86/0.16) | -0.32<br>(-1.44/0.80) | | -0.72<br>(-1.45/0.01) |
| Men | | | 1.74***<br>(1.04/2.44) | 1.6***<br>(0.95/2.25) |
| $R^2$ / Adjusted $R^2$ | 0.20 / 0.14 | 0.01 / -0.06 | 0.48 / 0.46 | 0.60 / 0.55 |
| AIC | 96.47 | 102.73 | 81.48 | 77.97 |
| F Statistic | 3.36* | 0.19 | 25.92*** | 12.77*** |

Reference: "low" levels and "women". p-values: (*) <0.05, (**) <0.01, (***) <0.001

**Supplementary Table 7. Multiple Linear Regression Models Predicting Physio PC2 Using Behavioral Variables.** Regression coefficients (95% CI) are reported for each category of independent variables. Model fit statistics include $R^2$, adjusted $R^2$, AIC and the F statistic.

| | Models | | | | |
|---|---|---|---|---|---|
| | **Daily Steps** | **Total MET** | **Vigorous MET** | **Alcohol** | **Marijuana** |
| Intercept | -0.14<br>(-0.81/0.53) | -0.41<br>(-1.22/0.40) | -0.46<br>(-1.27/0.35) | -0.01<br>(-0.51/0.49) | -0.04<br>(-0.52/0.44) |
| > 10000 steps/day | 0.26<br>(-0.65/1.17) | | | | |
| High METs levels | | 0.59<br>(-0.39/1.56) | 0.66<br>(-0.31/1.62) | | |
| High substance use | | | | 0.05<br>(-1.18/1.28) | 0.36<br>(-1.15/1.88) |
| $R^2$ / Adjusted $R^2$ | 0.01 / -0.02 | 0.05 / 0.02 | 0.07 / 0.03 | 0.00 / -0.04 | 0.01 / -0.03 |
| AIC | 100.78 | 99.56 | 99.13 | 101.14 | 100.89 |
| F Statistic | 0.35 | 1.52 | 1.95 | 0.01 | 0.24 |

Reference: "<10000 steps/day" and "low" levels. p-values: (*) <0.05, (**) <0.01, (***) <0.001

**Supplementary Table 8. Multiple Linear Regression Models Predicting Reaction Time.** Regression coefficients (95% CI) are reported for each category of independent variables, comparing main effects of physio PC1 and sex, with the combined variable. Model fit statistics include $R^2$, adjusted $R^2$, AIC and the F statistic.

| | **Models** | |
|---|---|---|
| | **Physio PC1 + Sex** | **Combined (Physio PC1 – Sex)** [a] |
| Intercept | 311.75*** (289.95/333.55) | 324.60*** (300.73/348.47) |
| Medium PC1/2 | -29.36* (-58.32/-0.40) | |
| High PC1/2 | -37.29** (-63.05/-11.53) | |
| Men | -24.19* (-47.98/-0.40) | |
| High PC1 - Men | | -67.60*** (-98.03/-37.17) |
| Low PC1 - Men | | -61.10*** (-89.52/-32.68) |
| High PC1 - Women | | -61.40*** (-95.16/-27.64) |
| $R^2$ / Adjusted $R^2$ | 0.42 / 0.35 | 0.49 / 0.43 |
| AIC | 290.39 | 286.26 |
| F Statistic | 6.16** | 8.35*** |

Reference: "low" levels and "women". p-values: (*) <0.05, (**) <0.01, (***) <0.001.
[a] Residual analysis: model does not meet the assumption of homoscedasticity.

**Supplementary Table 9. Multiple Linear Regression Models Predicting Reaction Time Using Plasma and Physio PCs and Sex Variables.** Regression coefficients (95% CI) are reported for each category of independent variables. Model fit statistics include $R^2$, adjusted $R^2$, AIC and the F statistic.

| | Models | | | | |
|---|---|---|---|---|---|
| | **Plasma PC1** | **Plasma PC2** | **Physio PC1** | **Physio PC2** | **Sex** |
| Intercept | 282.78*** (259.37/306.19) | 257.56*** (234.44/280.67) | 301.00*** (280.82/321.18) | 284.33*** (260.95/307.72) | 293.90*** (273.72/314.08) |
| Medium PC1/2 | -6.56 (-39.66/26.55) | 12.78 (-19.91/45.47) | -40.11** (-68.65/-11.57) | -10.33 (-43.41/22.74) | |
| High PC1/2 | -22.28 (-53.25/8.69) | 26.28 (-4.30/56.86) | -42.67** (-69.36/-15.97) | -23.33 (-54.27/7.60) | |
| Men | | | | | -33.00* (-57.71/-8.29) |
| $R^2$ / Adjusted $R^2$ | 0.08 / 0.01 | 0.10 / 0.04 | 0.32 / 0.27 | 0.08 / 0.02 | 0.21 / 0.18 |
| AIC | 301.96 | 301.20 | 293.05 | 301.90 | 295.39 |
| F Statistic | 1.19 | 1.57 | 6.27** | 1.22 | 7.48* |

Reference: "low" levels and "women". p-values: (*) <0.05, (**) <0.01, (***) <0.001

**Supplementary Table 10. Multiple Linear Regression Models Predicting Reaction Time Using Behavioral Variables.** Regression coefficients (95% CI) are reported for each category of independent variables. Model fit statistics include $R^2$, adjusted $R^2$, AIC and the F statistic.

| | Models | | | | |
|---|---|---|---|---|---|
| | **Daily Steps** | **Total MET** | **Vigorous MET** | **Alcohol** | **Marijuana** |
| Intercept | 276.43*** (257.38/295.47) | 277.89*** (254.11/301.67) | 278.78*** (255.05/302.51) | 270.40*** (256.11/284.69) | 271.81*** (257.99/285.64) |
| > 10000 steps/day | -8.49 (-34.57/17.59) | | | | |
| High METs levels | | -8.56 (-36.98/19.87) | -9.83 (-38.19/18.54) | | |
| High substance use | | | | 9.00 (-26.01/44.01) | 0.85 (-42.86/44.56) |
| $R^2$ / Adjusted $R^2$ | 0.02 / -0.02 | 0.01 / -0.02 | 0.02 / -0.02 | 0.01 / -0.03 | 0.00 / -0.04 |
| AIC | 302.02 | 302.09 | 301.96 | 302.20 | 302.49 |
| F Statistic | 0.44 | 0.38 | 0.50 | 0.28 | 0.00 |

Reference: “<10000 steps/day” and "low" levels. p-values: (*) <0.05, (**) <0.01, (***) <0.001

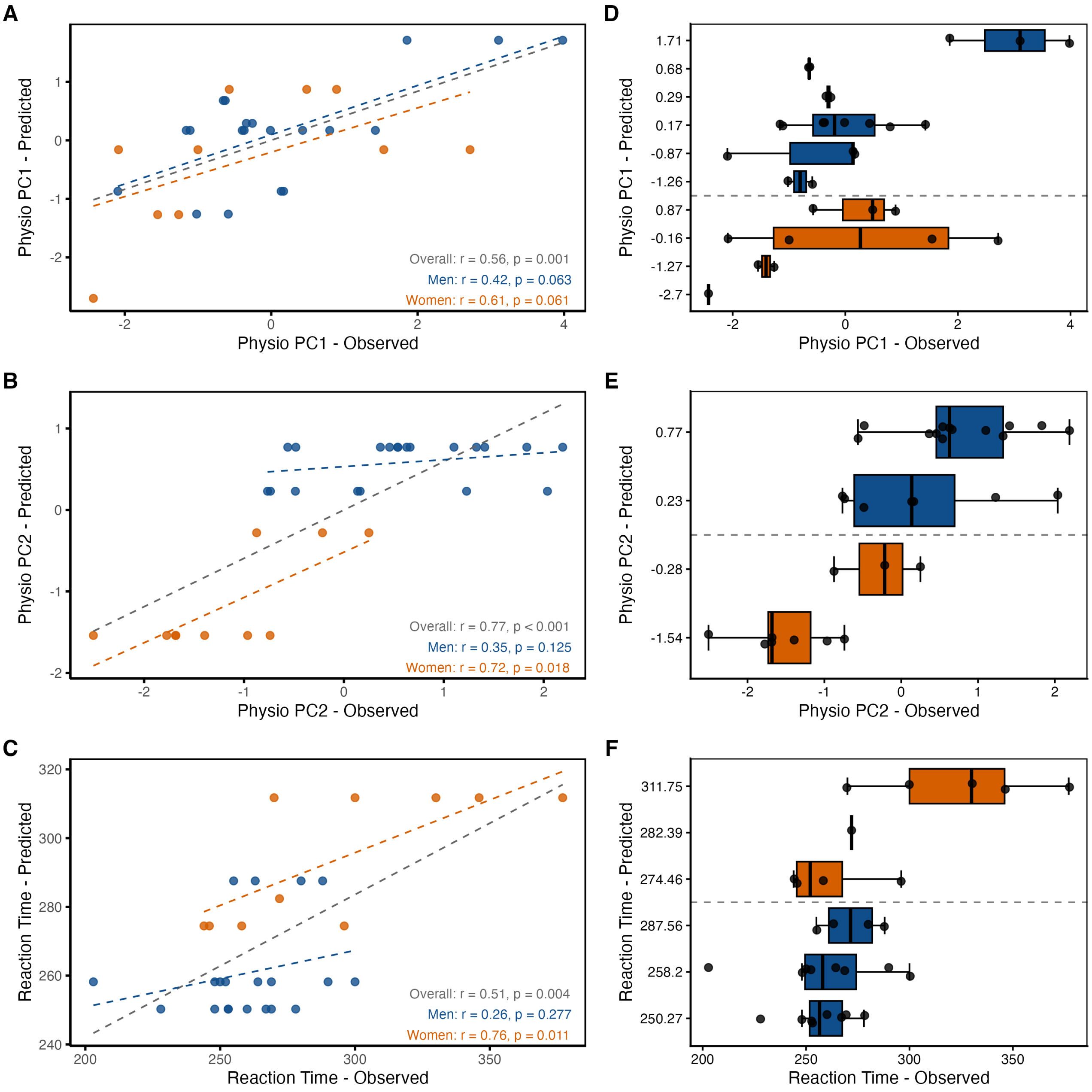


**Supplementary Figure 1: Association between observed and predicted values.** (**A-C**) Representation of overall (n = 30), men (blue, n = 20), and women (orange, n = 10) performance groups for (**A**) Physio PC1, (**B**) Physio PC2 and (**C**) RT models. Spearman's coefficients (r) and p-values were determinated. (**D–E**) The application of the models showed the segregation of the sample into groups of men (blue) and women (orange), according to the categorization of the independent variables.
